## Extended Data for "Distinct mechanisms for TMPRSS2 expression explain organ-specific inhibition of SARS-CoV-2 infection by enzalutamide"

### Extended Data Figure 1

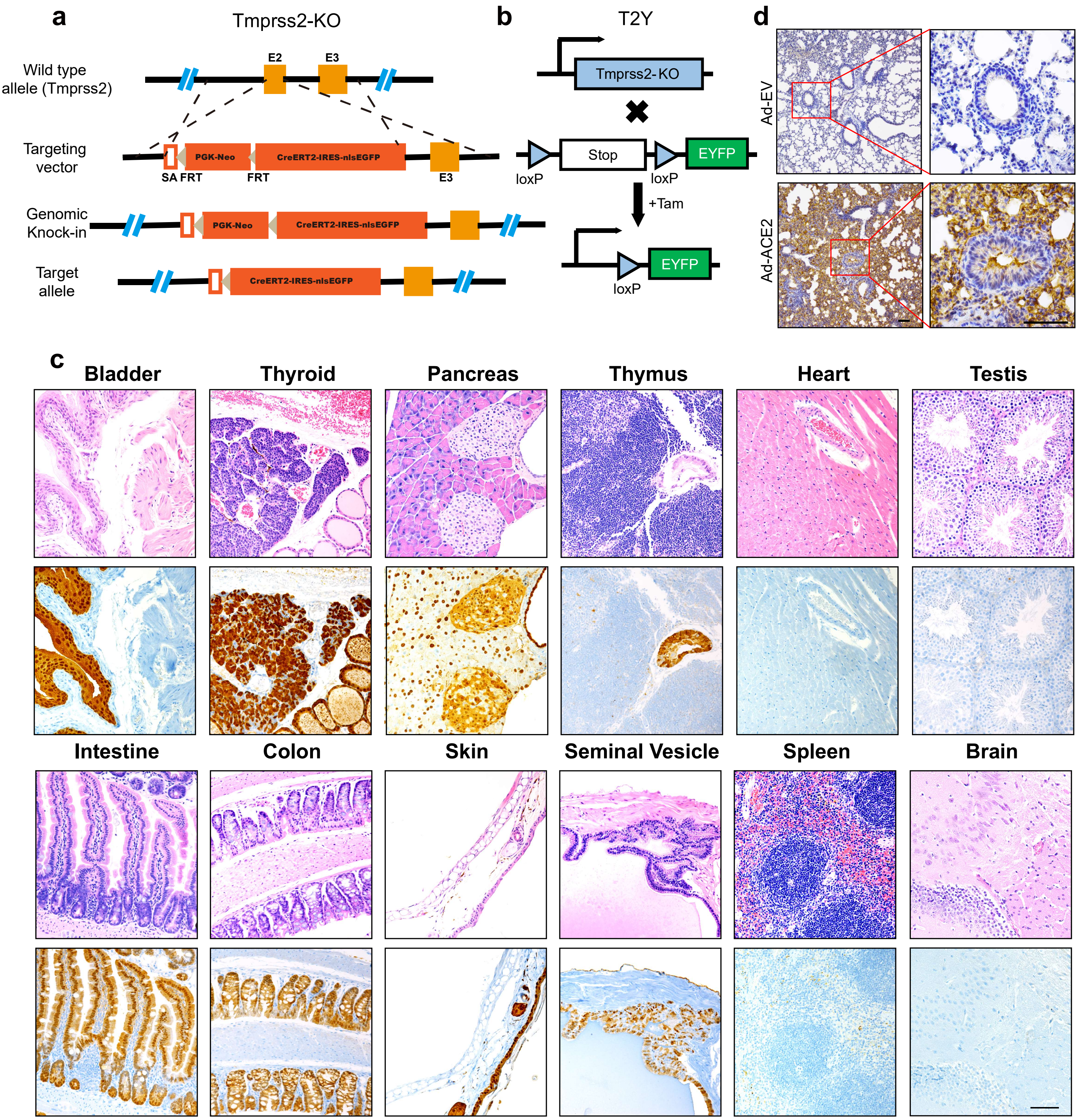

**Extended Data Figure 1 Identification of Tmprss2-positive cells in multiple mouse organs. (a)** Construction strategy for Tmprss2-KO mice. **(b)** Breeding strategy for the generation of T2Y mice. **(c)** YFP IHC staining for multiple organs of T2Y mice with tamoxifen gavage. **(d)** FLAG IHC staining in the lungs of WT transduced with Ad-EV (top) or Ad-ACE2 transduction (bottom), samples were collected 5 days post transduction.

### Extended Data Figure 2

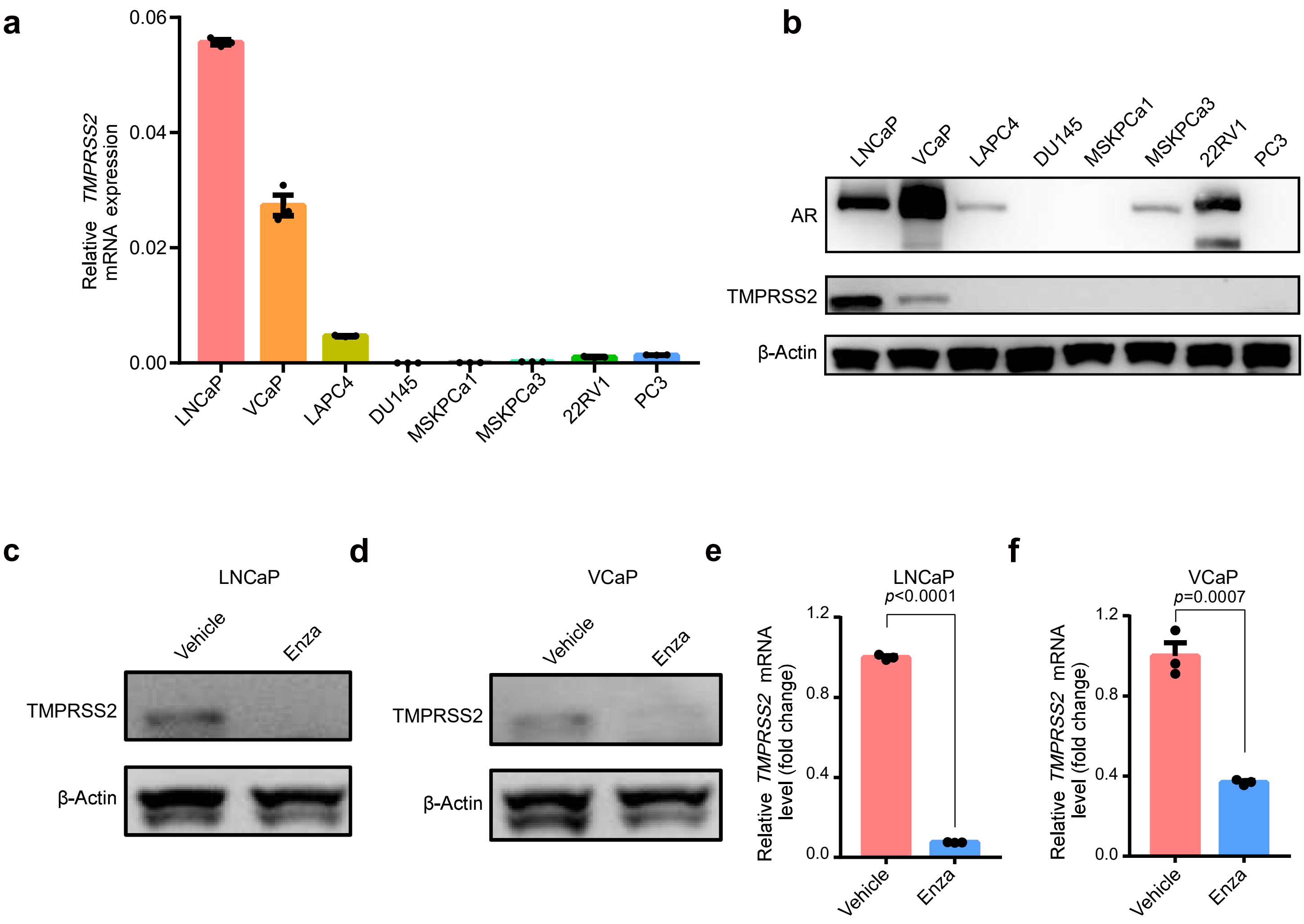

**Extended Data Figure 2 TMPRSS2 expression is reduced by enzalutamide in LNCaP and VCaP cells. (a)** qRT-PCR analysis of *TMPRSS2* mRNA expression in multiple prostate cancer cell lines and two prostate cancer organoid lines. **(b)** Western blotting analysis of TMPRSS2 and AR expression in multiple prostate cancer cell lines and two prostate cancer organoid lines. **(c)** Western blotting analysis of TMPRSS2 expression in LNCaP cells treated with vehicle under normal FBS condition or enzalutamide treatment under charcoal stripped FBS condition. **(d)** Western blotting analysis of TMPRSS2 expression in VCaP cells treated with vehicle under normal FBS condition or enzalutamide treatment under charcoal stripped FBS condition. **(e)** qRT-PCR analysis of *TMPRSS2* mRNA expression in LNCaP cells treated with vehicle under normal FBS condition or enzalutamide treatment under charcoal stripped FBS condition (two-tailed t-test, mean ± SEM, n=3). **(f)** qRT-PCR analysis of *TMPRSS2* mRNA expression in VCaP cells treated with vehicle under normal FBS condition or enzalutamide treatment under charcoal stripped FBS condition (two-tailed t-test, mean ± SEM, n=3).

#### Extended Data Figure 3

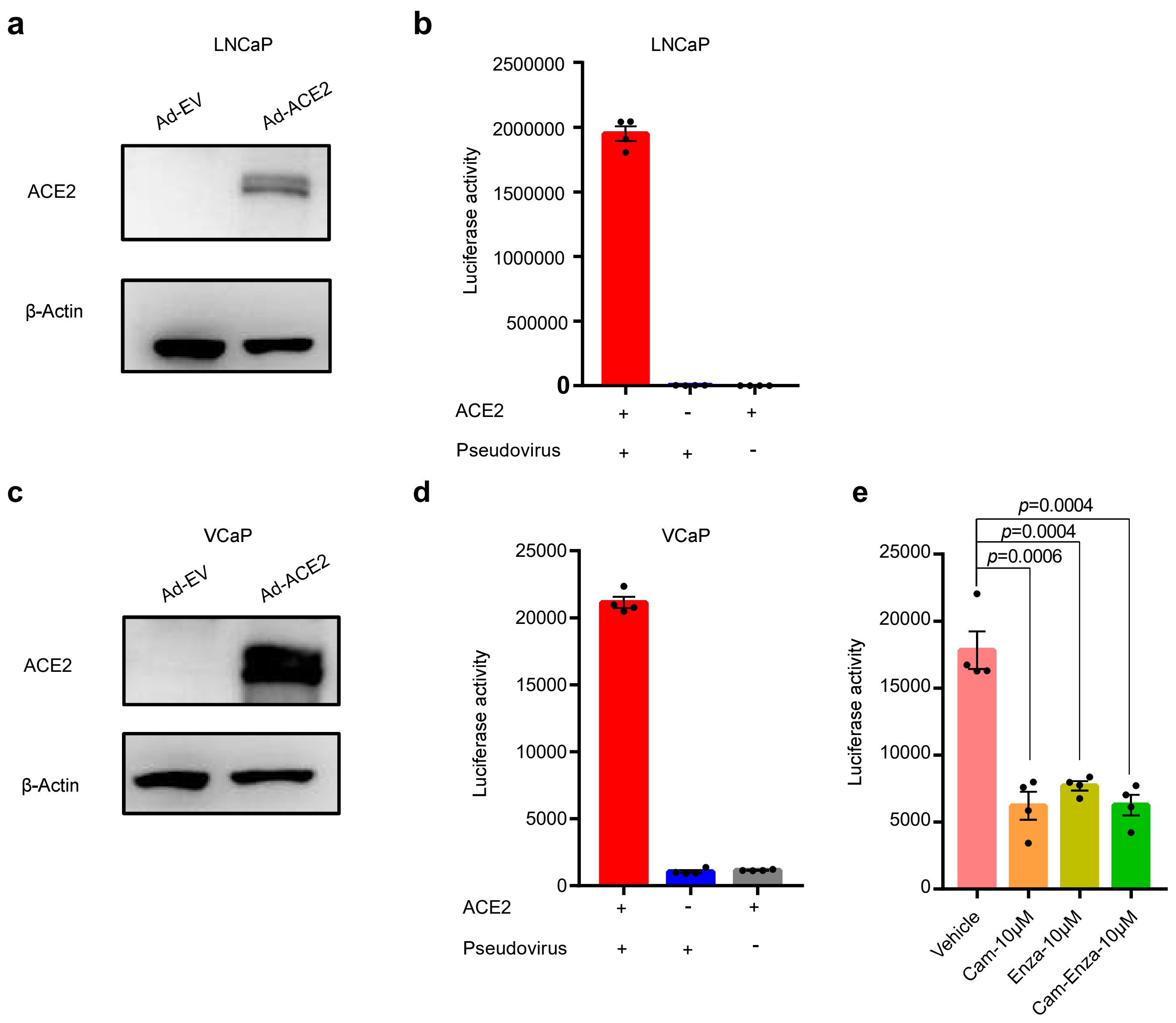

**Extended Data Figure 3 Enzalutamide significantly reduces infection of VCaP cell with SARS-CoV-2-S. (a)** Western blotting analysis of ACE2 and β-Actin expression in LNCaP cells with empty vector adenovirus transduction or ACE2 adenovirus transduction. **(b)** SARS-CoV-2-S-driven entry into LNCaP cells with or without Ad-ACE2 transduction. Luciferase activity was measured 48 hours post SARS-CoV-2-S infection. **(c)** Western blotting analysis of ACE2 and β-Actin expression in VCaP cells with empty vector adenovirus transduction or ACE2 adenovirus transduction. **(d)** SARS-CoV-2-S-driven entry into VCaP cells with or without Ad-ACE2 transduction, luciferase activity was measured 48 hours post SARS-CoV-2-S infection. **(e)** SARS-CoV-2-S-driven entry into VCaP cells with Ad-ACE2 transduction treated with vehicle, 10 μM camostat mesylate, 10 μM enzalutamide and 10 μM camostat mesylate/enzalutamide. Luciferase activity was measured 48 hours post SARS-CoV-2-S infection (two-tailed t-test, mean ± SEM, n=4).

**Extended Data Figure 4**

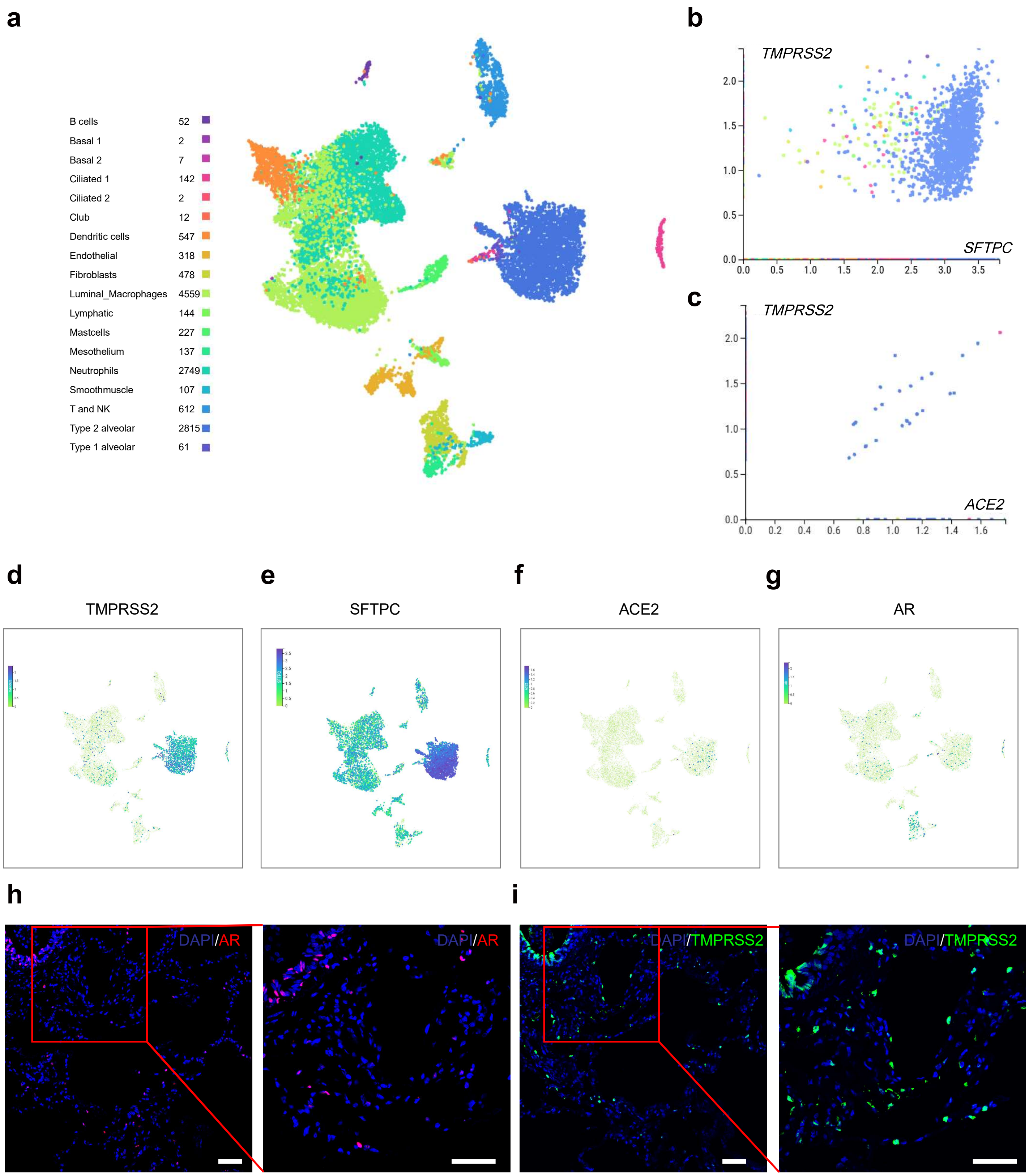

**Extended Data Figure 4 AR-positive and TMPRSS2-positive cells are verified in human lungs. (a)** UMAP plot displaying cell clusters of human airway epithelial cells from a public dataset using COVID-19 Cell Atlas. **(b)** *TMPRSS2* and *SFTPC* mRNA expression across multiple cell types. **(c)** *TMPRSS2* and *ACE2* mRNA expression across multiple cell types. **(d-g)** *TMPRSS2* **(d)**, *SFTPC* **(e)**, *ACE2* **(f)** and *AR* **(g)** mRNA expression levels on UMAP plots across multiple cell types. **(h-i)** Immunofluorescence staining of AR **(h)** and TMPRSS2 **(i)** in human lungs using adjacent sections. Scale bars represent 50 μm.

### Extended Data Figure 5

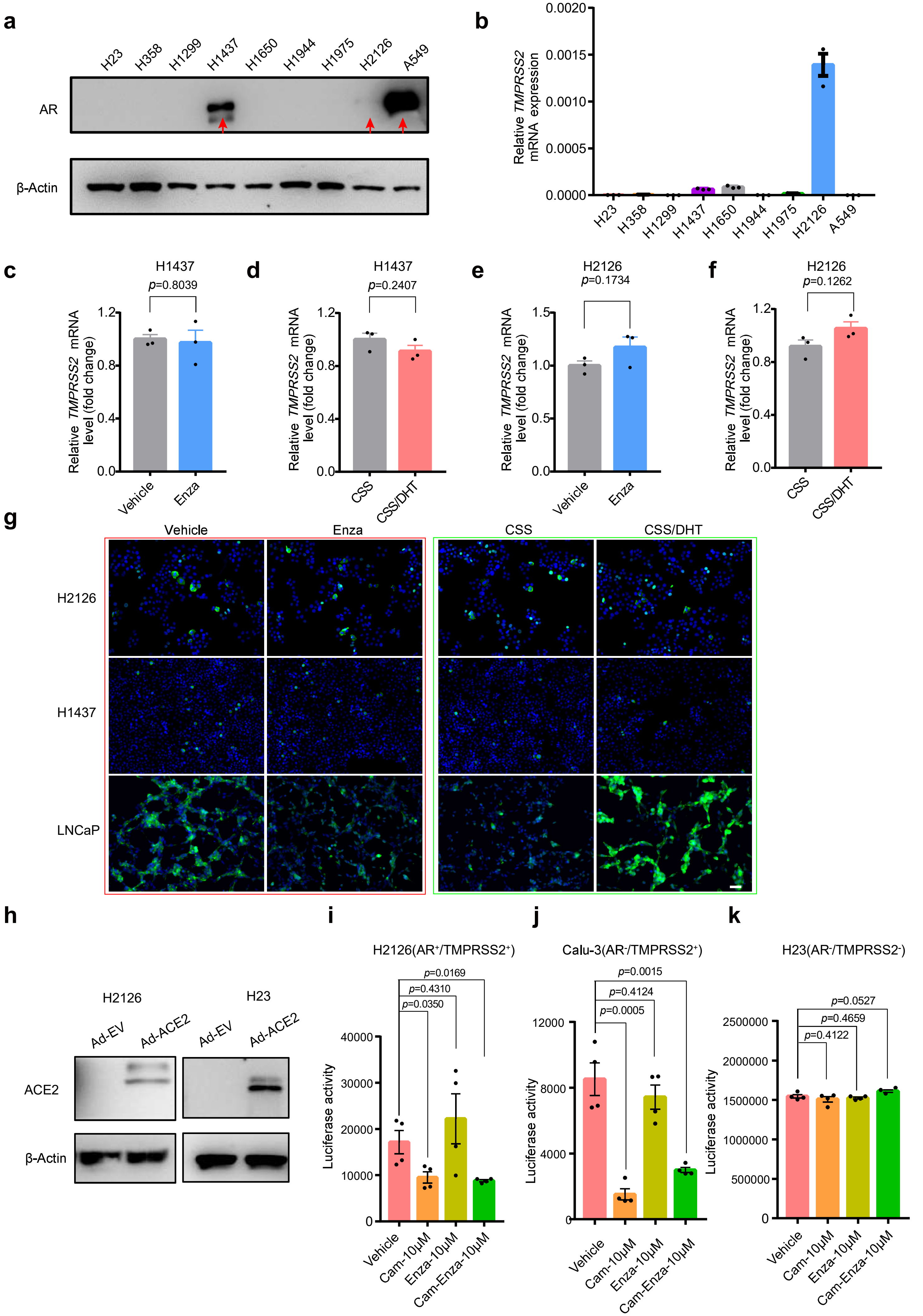

**Extended Data Figure 5 Enzalutamide does not prevent SARS-CoV-2-S-driven entry into lung cells.** **(a)** Western blotting analysis of AR expression in multiple lung cancer cells. **(b)** qRT-PCR analysis of *TMPRSS2* mRNA expression in multiple lung cancer cells. **(c)** qRT-PCR analysis of *TMPRSS2* mRNA expression in H1437 cells with vehicle or enzalutamide treatment. **(d)** qRT-PCR analysis of *TMPRSS2* mRNA expression in H1437 cells with or without DHT treatment under charcoal stripped FBS condition. **(e)** qRT-PCR analysis of *TMPRSS2* mRNA expression in H2126 cells with vehicle or enzalutamide treatment. **(f)** qRT-PCR analysis of *TMPRSS2* mRNA expression in H2126 cells with or without DHT treatment under charcoal stripped FBS condition. **(g)** Immunofluorescence staining of TMPRSS2 in H2126 (top), H1437 (middle) and LNCaP (bottom) cells under multiple treatments condition. **(h)** Western blotting analysis of ACE2 and β-Actin expression in H2126 (left) and H23 (right) cells with empty vector adenovirus transduction or ACE2 adenovirus transduction respectively. **(i-k)** SARS-CoV-2-S-driven entry into H2126 **(i)**, Calu-3 **(j)** and H23 **(k)** cells transduced with Ad-ACE2 and treated with vehicle, 10 μM camostat mesylate, 10 μM enzalutamide and 10 μM camostat mesylate/enzalutamide. Luciferase activity was measured 48 hours post SARS-CoV-2-S infection (two-tailed t-test, mean ± SEM, n=4). Scale bars represent 50 μm.

**Extended Data Figure 6**

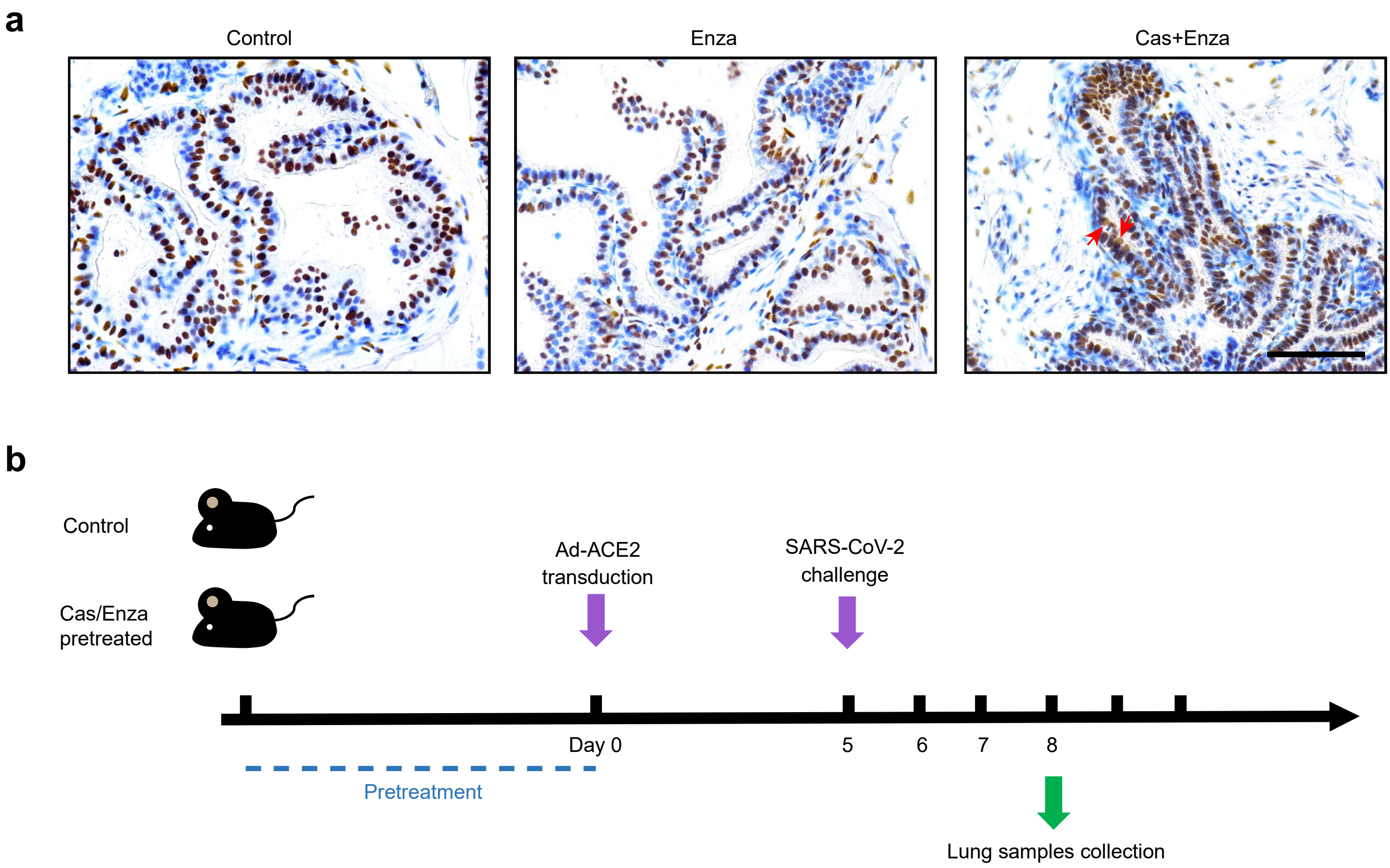

**Extended Data Figure 6 Ad-ACE2-transduced mouse models are employed to assess the efficacy of enzalutamide *in vivo*. (a)** IHC staining for AR in prostates of wild type mice, enzalutamide-treated mice and enzalutamide-treated castrated mice respectively. Red arrows indicate blocking of nuclear translocation of AR in prostates of enzalutamide treated castrated mice. **(b)** Schematic strategy for Ad-ACE2-transduced mouse to evaluate the therapeutic efficacy of enzalutamide.

### Extended Data Figure 7

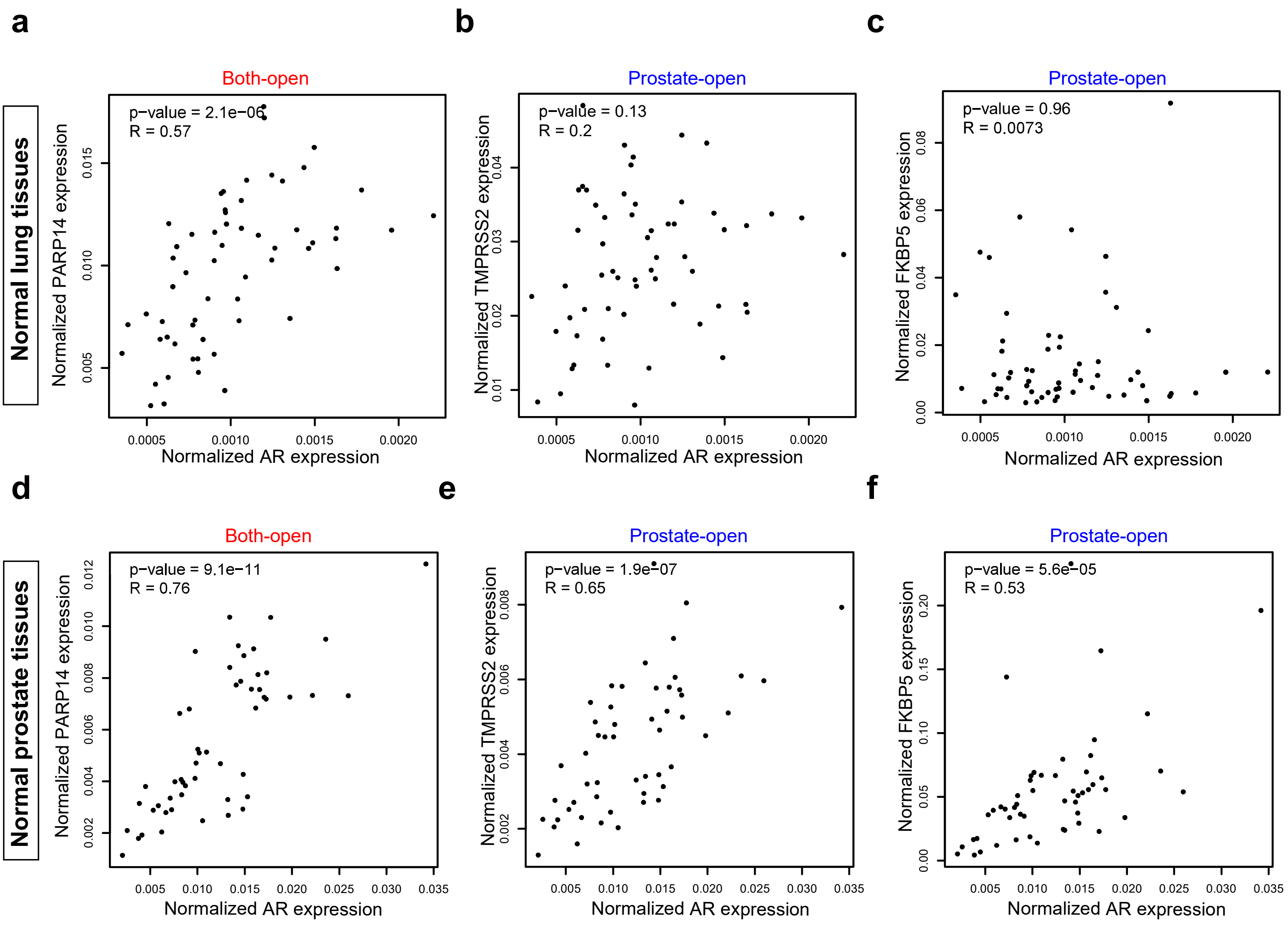

**Extended Data Figure 7 Correlation analysis of mRNA levels of AR and TMPRSS2 in human lung tissues and prostate tissues. (a-c)** Correlation analysis for normalized expression of AR and a both-open gene *PARP14* (left) **(a)**, two prostate-open genes *TMPRSS2* (middle) **(b)** and *FKBP5* (right) respectively in normal lung tissues using TCGA datasets **(c)**. **(d-f)** Correlation analysis for normalized expression of *AR* and a both-open gene *PARP14* (left) **(d)**, two prostate-open genes *TMPRSS2* (middle) **(e)** and *FKBP5* (right) respectively in normal prostate tissues using TCGA datasets **(f)**.
